# Antibody-dependent enhancement representing *in vitro* infective progeny virus titer correlates with the viremia level in dengue patients

**DOI:** 10.1101/2020.11.20.392357

**Authors:** Atsushi Yamanaka, Hisham Ahmed Imad, Weerapong Phumratanaprapin, Juthamas Phadungsombat, Eiji Konishi, Tatsuo Shioda

**Author notes:** Corresponding author: Atsushi Yamanaka, Ph.D. Mahidol-Osaka Center for Infectious Diseases, Faculty of Tropical Medicine, Mahidol University, 420/6 Ratchawithi Road, Ratchathewi, Bangkok 10400, Thailand.

## Abstract

Dengue virus (DENV) distributes throughout tropical and subtropical countries and causes dengue fever (DF) and dengue hemorrhagic fever in humans. Some DF patients suddenly develop severe symptoms after the defervescent period. Although the pathogenic mechanism of the severe symptoms has not been fully elucidated, the viremia level in the early phase has been shown to correlate with the disease severity. One of the hypotheses is that a phenomenon called antibody-dependent enhancement (ADE) of infection leads to a high level of viremia. To examine the plausibility of this hypothesis, we examined the relationship between *in vitro* ADE activity and *in vivo* viral load quantity in six patients with dengue diseases. An autologous DENV strain was isolated from each of the six patients. Blood samples were then collected at multiple time points between the acute and defervescent phases, and the balance between neutralizing and enhancing activities against the autologous and prototype viruses was examined. As the antibody levels against DENV were rapidly increased, ADE activity was decreased over time or partially maintained against some viruses at low serum dilution. In addition, positive correlations were observed between ADE activity representing *in vitro* progeny virus production and viremia levels in patient plasma samples. Therefore, the measurement of ADE activity in dengue-seropositive samples may help to predict the impact of viral load in the subsequent DENV infection.

**IMPORTANCE:** It has not been fully elucidated how the phenomenon of antibody-dependent enhancement (ADE) affects the pathogenesis of severe dengue diseases, although high viremia levels have been epidemiologically demonstrated to be associated with the disease severity. Here, we show that ADE in the acute-phase patient sera exhibited significantly different activities against autologous and lab strains than ADE in the defervescent-phase sera. Further, the enhancement of progeny virus production activity, which is one of the factors to evaluate ADE *in vitro*, was significantly correlated with the levels of viral load in the patient blood circulation. This suggests that measurement of the *in vitro* enhancing progeny virus titers might be used to predict the impact of *in vivo* DENV viremia level. Our present findings could contribute to a method to forecast disease severity for seropositive populations who would be at risk of developing severe disease in the event of heterotypic DENV infection.

## INTRODUCTION

Dengue virus (DENV), belonging to the family *Flaviviridae*, genus *Flavivirus*, is distributed throughout tropical and subtropical areas of the world (1). DENV is transmitted by *Aedes* mosquito species and causes dengue fever (DF) and severe dengue in humans. Approximately 3.9 billion people are under the risk of infection (2). An estimated 390 million people are infected with DENV annually, and 100 million of these individuals show clinical symptoms (3). Therefore, dengue is one of the most important mosquito-borne viral diseases worldwide, and it should be controlled to the greatest extent possible.

The four serotypes of DENV (DENV-1, DENV-2, DENV-3 and DENV-4) are genetically distinct, and there is complicated immunological cross-reactivity among them (4). Secondary heterotypic infection has been epidemiologically demonstrated to give rise to the severe forms—namely, dengue hemorrhagic fever (DHF) and dengue shock syndrome (DSS) (5). Patients showing dengue with warning signs have a risk of developing disease severity, with the emergence of severity usually occurring around the defervescence phase, beginning at days 3-7 of illness (6, 7). The mortality rate of cases with DSS is much higher than that of cases without DSS (8). Although a mechanism associated with the severity and a surrogate marker predicting the deterioration have not been fully identified yet, high levels of viremia have been shown to be related to disease severity (6, 9, 10). Antibody-dependent enhancement of infection (ADE) has been proposed as one of the pathogenic mechanisms that increases the viremia level; in the case of ADE the increase occurs by viral internalization via Fc gamma receptors (11). Recently, a potential relationship between ADE and human disease severity in DENV infection has been reported (12). However, it is still unclear whether *in vitro* ADE can be used for the prediction of subsequent clinical outcomes.

Enhancing antibodies (EAbs), which exclusively play a role of the ADE phenomenon (13), may be associated with an increase in the viremia level in DENV infection (14, 15). In contrast, neutralizing antibodies (NAbs) have a biological function to decrease the viremia level to protect the host from DENV infection (16), while most NAbs show ADE activity at subneutralizing doses (17). A DENV-immune serum (polyclonal form) could be represented with a cocktail of these intricate functional antibodies (monoclonal EAbs and NAbs) (18), and could be introduced by one of three routes: (i) DENV infection, (ii) maternal antibody from a DENV-seropositive mother and (iii) other flavivirus infection. Therefore, the balance activity between EAbs and NAbs might be critical to control the outcome (protection or pathogenesis). We previously developed a simple method to detect the balance between the enhancing and neutralizing activities (19), and demonstrated that mouse monoclonal EAbs and NAbs competed over the neutralizing activities *in vitro* (18, 20). Specifically, the neutralizing activity of an NAb was reduced in the presence of a sufficient level of an EAb, suggesting that the relative capacity for neutralization might be easily affected by the balance between NAbs and EAbs.

In the present study, we evaluated the balance between neutralizing and enhancing activities in sera collected from dengue patients at multiple time points between the acute and defervescent phases. The six autologous viruses isolated from the respective patients were used as assay antigens, allowing us to examine the balance antibody assay with autologous combinations between patient sera and virus antigens. We also measured the number of viral RNA copies in plasma samples collected at multiple time points, and revealed a correlation between the *in vitro* ADE activity and *in vivo* viral load quantity.

## RESULTS

### Balance between the neutralizing and enhancing activities against autologous viruses

Six hospitalized adult patients, who were enrolled in the previous study (21), were recruited into the present study, and an autologous DENV clinical strain was successfully isolated from each of them. The demographic information (infecting serotype, diagnosis [DF or DHF], ID number, days after fever onset and sample collection period [h] after the hospitalization) is shown in Fig. 1A. Serum samples, which were collected at several time points between the acute and defervescent phases, were subjected to an antibody assay to determine the balance between neutralizing and enhancing activities (NAb/EAb-balance assay) using each autologous virus. Fig. 1B shows that the serum dilution displaying the highest number of infected cells clearly shifted from low to high over time after the first sampling in four of the patients (#44, #57, #12 and #49). These results indicate that their antibody levels rapidly increased by 16~256 fold during the early disease stage. Similar dose-dependent patterns were observed in patients #44, #57 and #49, even though these patients had different infecting serotypes and diagnoses. On the other hand, patients #14 and #46 did not show a remarkable shift during the observation period. As patient #14 was classified as primary infection by IgG/IgM immunochromatography tests in the previous study (21), no functional antibody activity was observed. Although patient #46 showed strong neutralization with a wide range of serum dilutions (1:10~1:2560), this patient was diagnosed with DHF with severe clinical symptoms.

**FIG 1.**
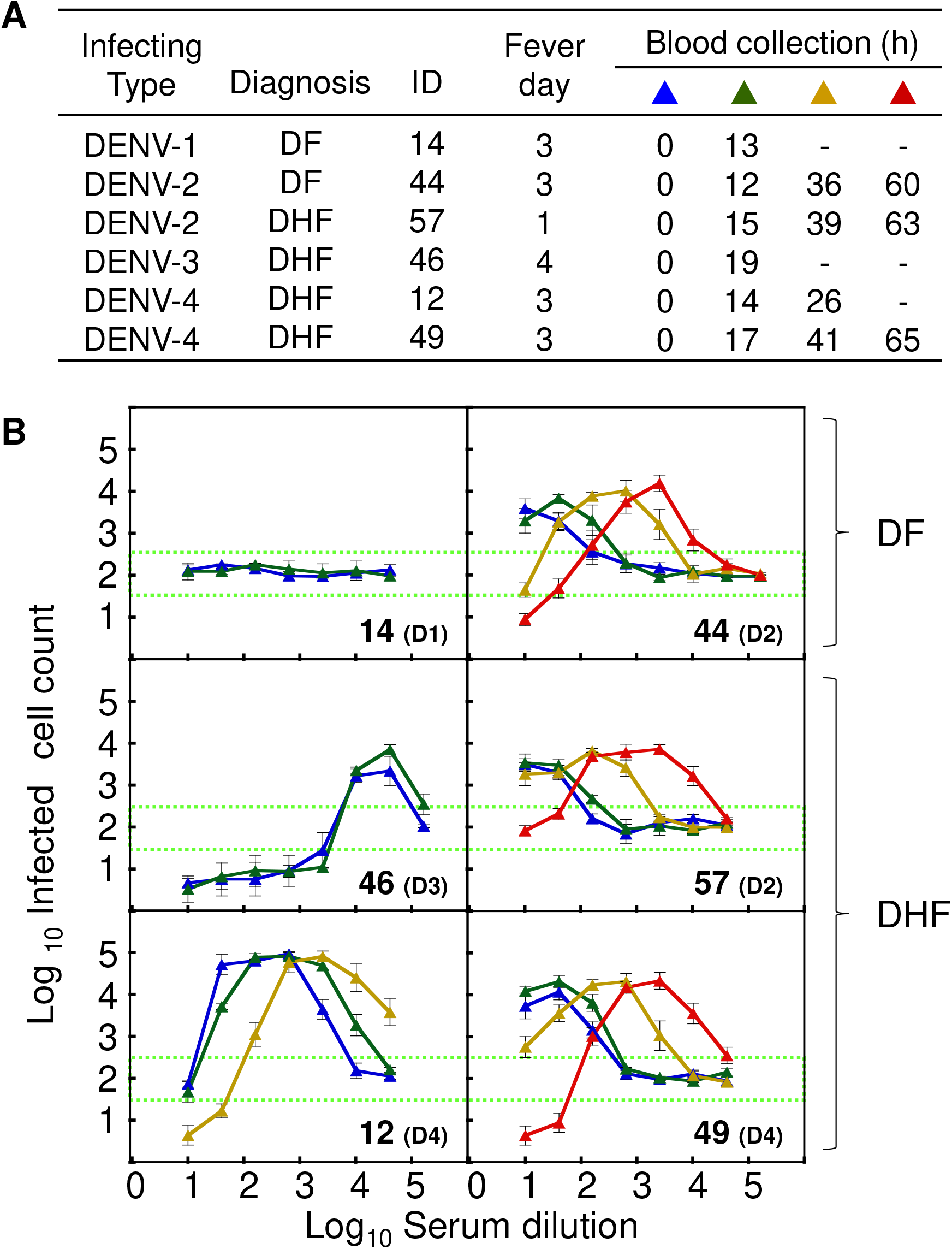
Balance between neutralizing and enhancing activities against autologous virus. (A) Demographic data for the six dengue patients enrolled in this study. Blood samples were collected on the day of hospitalization and the following time points at 12-24 h intervals until a maximum 65 h after hospitalization. (B) NAb/EAb balance activity against the autologous virus. The NAb/EAb-balance assay was performed using a combination of autologous virus and serum. The *blue, green, yellow* and *red* triangles, corresponding to Fig. 1A, indicate the hours after hospitalization. The infecting serotype is indicated in parentheses (D1: DENV-1; D2: DENV-2; D3: DENV-3; and D4: DENV-4). The abscissa indicates the serum dilutions and the ordinate shows the numbers of infected cells (both expressed as log10). Each data point represents the average of two separate assays; error bars indicate the SDs. Dotted lines indicate the mean numbers of infected cells plus or minus three times the SD (mean ± 3SD) calculated from eight negative controls.

### Balance between neutralizing and enhancing activities against four prototype viruses

Dose-dependent neutralizing/enhancing activity curves were obtained against four prototype laboratory strains (DENV-1: Mochizuki; DENV-2: NGC; DENV-3: H87; and DENV-4: H241). As described in the above assays using the autologous viruses (Fig. 1B), increases in the antibody levels against four prototype DENVs were observed in four patients (#44, #57, #12 and #49) (Fig. 2). Although patients #12 and #49 were currently infected with DENV-4, they displayed higher levels of cross-neutralization against heterologous DENV-2 and/or DENV-1. Similarly, patient #57, who was currently infected with DENV-2, showed cross-neutralization against DENV-4. These results suggest that our assay system may be able to reveal the previous infection history in dengue patients by comparing their cross-neutralizing activities. On the other hand, patients #14 and #46 did not show any remarkable antibody shift against the four prototype viruses. Patient #46 displayed strong neutralization against all serotypes, while patient #14 showed enhancing activities against DENV-1 and DENV-3 at 1:10 serum dilution. Based on these findings, it seems likely that a history of previous exposure to and infection by DENV affected the antibody response patterns in our patients.

**FIG 2.**
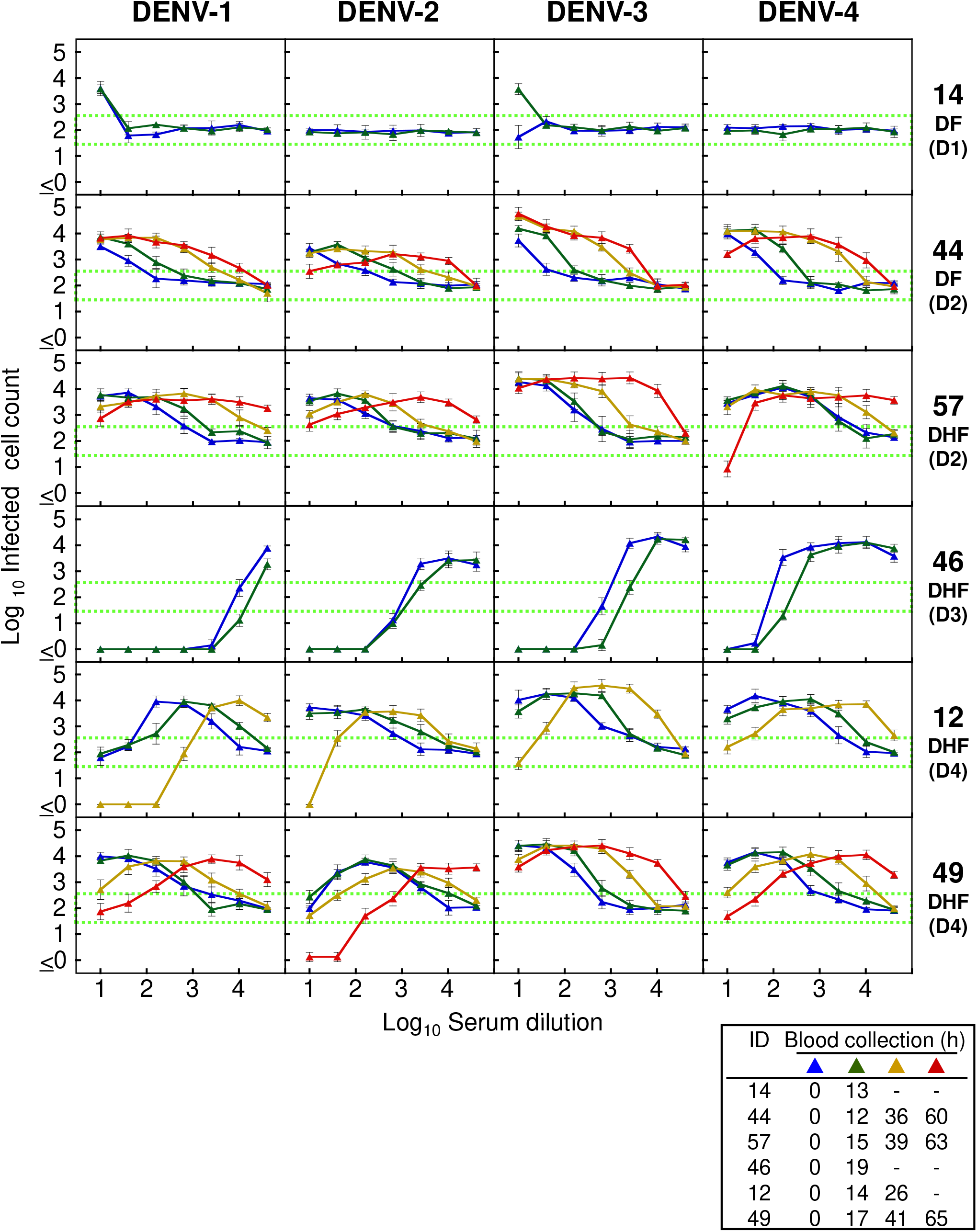
Balance between neutralizing and enhancing activities against four prototype viruses. The NAb/EAb-balance assay was performed using a combination of serum and four prototype viruses (DENV-1: Mochizuki strain; DENV-2: NGC strain; DENV-3: H87 strain; and DENV-4: H241 strain). The *blue, green,yellow* and *red* triangles, corresponding to Fig. 1A, indicate the hours after hospitalization. The infecting serotype is indicated in parentheses (D1: DENV-1; D2: DENV-2; D3: DENV-3; and D4: DENV-4). The abscissa indicates the serum dilutions and the ordinate shows the numbers of infected cells (both expressed as log10). Each data point represents the average of two separate assays; error bars indicate the SDs. Dotted lines indicate the mean numbers of infected cells plus or minus three times the SD (mean ± 3SD) calculated from eight negative controls.

### Evaluation of progeny virus titers by NAb/EAb balance assay

To evaluate the levels of progeny virus secreted from the infected cells using the NAb/EAb balance assay system, patient #49 was selected as a representative patient with a range of neutralizing and enhancing activities (Figs. 1 and 2). Culture supernatants were harvested from the infected cells 24 h after the set-up of the NAb/EAb-balance assay, and were titrated on Vero cells. The dose-dependent patterns of progeny virus titers (Fig. 3A) were roughly similar to those from the infected-cell counts (Figs. 1 and 2), with minor differences at lower serum dilutions. Specifically, the defervescent sample (65 h) displayed a trend of neutralization against four prototype viruses in approximately 1:10-1:100 serum dilutions. Fold enhancements were calculated from progeny virus titers at 1:10 serum dilutions (Fig. 3A), and were plotted as the ordinate against time (h) as the abscissae (Fig. 3B). The time-course patterns tended to decline in parallel, except for DENV-2. The autologous virus showed the higher fold enhancement (>1000 fold on 0 h) than the other four prototype assay antigens. In contrast, no infective progeny virus was detected in DENV-2, suggesting that patient #49 had previously been infected with DENV-2, and possessed sufficient neutralizing antibody against DENV-2. Similar declining trends were observed in graphic data based on infected-cell counts as the ordinate (Supplemental Fig. S1).

**FIG 3.**
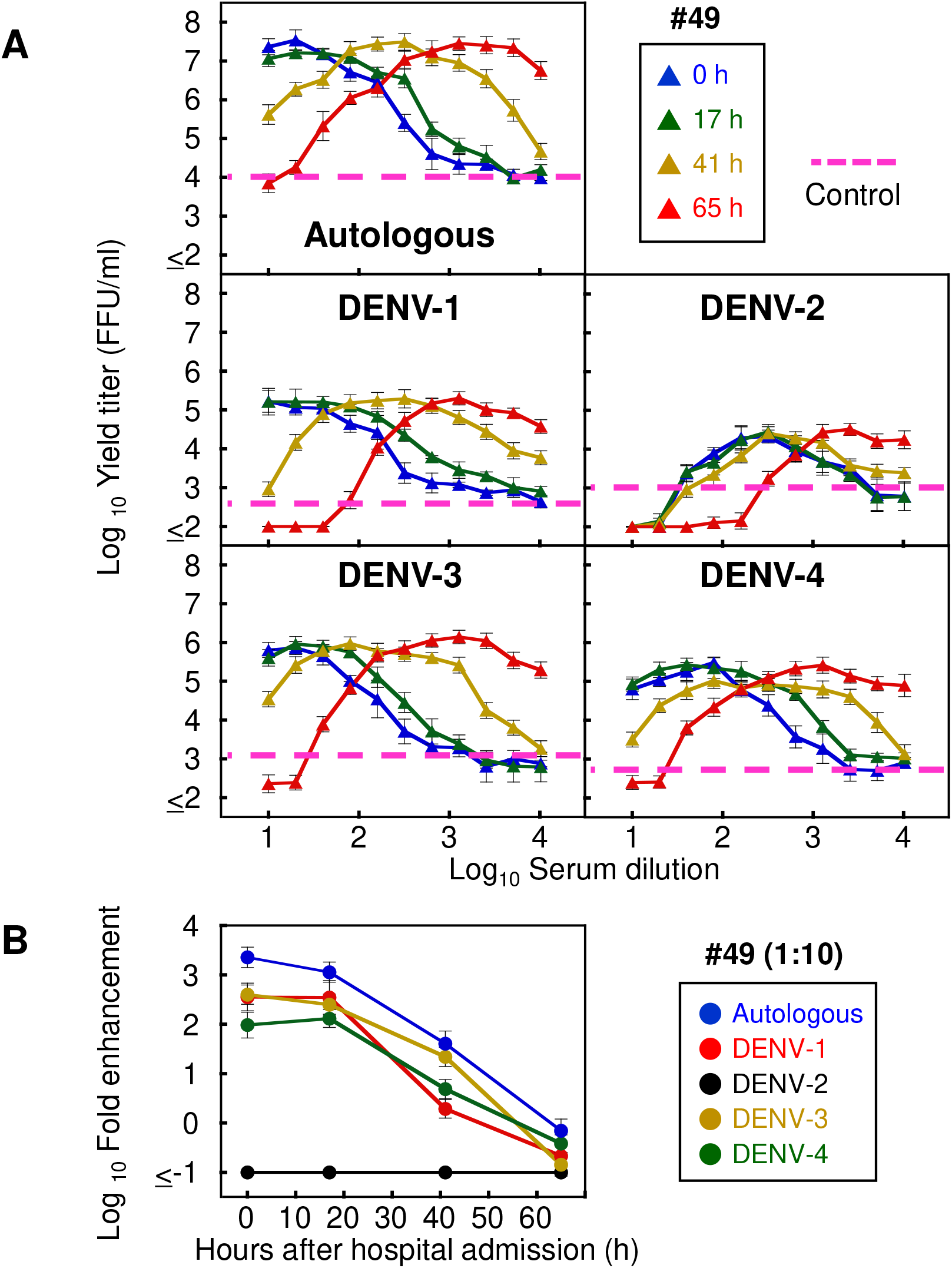
Progeny virus infectivity in the NAb/EAb balance assay. (A) Progeny virus titers obtained from the NAb/EAb-balance assay. The NAb/EAb-balance assay was conducted with a combination of 2-fold serial serum dilutions (starting from 1:10) in patient #49, and each DENV strain (the autologous, Mochizuki, NGC, H87 or H241 strain). Culture supernatants were harvested at 24 h after the mixture of serum, virus and K562 cells. The infective titers (FFU/ml) were determined on Vero cells. The *blue, green, yellow* and *red* triangles, corresponding to Fig. 1A, indicate the hours after hospitalization. The abscissa indicates the serum dilutions and the ordinate shows the progeny virus titers (both expressed as log10). *Dotted lines* indicate the mean progeny virus titers calculated from four negative (no serum) controls. (B) Decreasing trend of ADE at low serum dilution with time progression. Fold enhancement was calculated from the infective progeny virus titers obtained in Fig. 3A (specifying data at 1:10 serum dilution), and was expressed in log10 as the increase in the progeny virus titer relative to the negative control. The fold enhancement was plotted as the ordinate against time (h) as the abscissa.

### Relationship between the *in vitro* infective progeny virus and *in vivo* viral load

Using the same method described above, the infection-enhanced progeny virus titers were determined with the autologous combination (using 1:10 serum dilution) in the other five patients (#14, #44, #57, #46 and #12) (blue triangles in Fig. 4A). In addition, the quantity of viral RNA was directly determined from the corresponding clinical plasma samples (red circles in Fig. 4A). Both the ADE progeny virus titer (blue triangle) and viral RNA copy number (red circle) basically declined in parallel over time. Significant correlation was observed between the ADE progeny virus titers and the viral RNA copy numbers (r=0.69; *P*<0.05) (Fig. 4B). These results indicate that there might be positive correlations between (i) the *in vitro* progeny virus titers obtained from the NAb/EAb-balance assay and (ii) the *in vivo* viral quantity in the plasma. In contrast, no correlations were observed between the laboratory results (the intensity of ADE activity) and the clinical disease outcome (DF or DHF). Patients #57, #46, #12 and #49 deteriorated after the beginning of the defervescent phase (*pink shading* in Fig. 4A), but the levels of progeny virus titers and viral load in #46 (DHF patient) were not as high as those in other DHF patients. Likewise, #44 showed DF manifestations, but the levels of progeny virus titers and viral load were as high as those of DHF patients.

**FIG 4.**
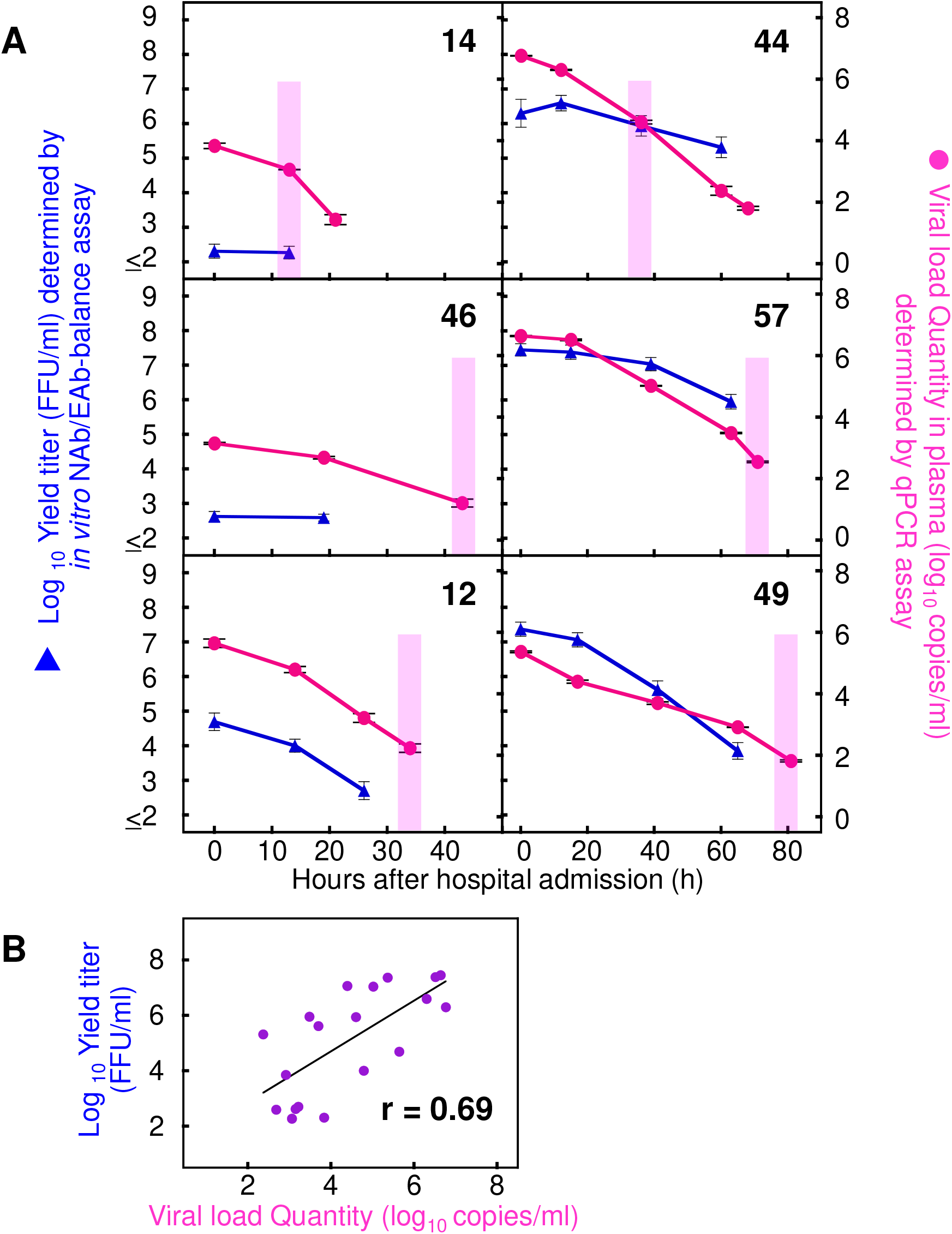
Relationship between *in vitro* ADE activity and *in vivo* viral load quantity. (A) The NAb/EAb-balance assay was conducted with a combination of serum diluted at 1:10 and the corresponding autologous virus. Culture supernatants were harvested at 24 h after the mixture of serum, virus and K562 cells. The infective titers were determined on Vero cells and plotted as the left ordinates (*blue triangles*: expressed as log10 FFU/ml). The number of viral RNA copies in plasma samples collected at same time points as well as one extra time point were determined by real time RT-PCR, and plotted as the right ordinates (*red circles:* expressed as log10 copies/ml). The abscissae indicate time (h) after the hospitalization. *Pink shading* indicates the beginning of defervescence on the clinical observation. (B) Correlation between the ADE progeny virus titers and the viral RNA copy numbers. The correlation coefficient (r) was estimated for the individual progeny virus titers and qPCR values obtained in Fig. 4A. The abscissa and ordinate indicate the qPCR values and the progeny virus titers, respectively. Linear regression lines and r values are presented in the panel.

## DISCUSSION

In the present study, to elucidate the relationship between antibody status and disease severity in dengue patients, we analyzed the balance between neutralizing and enhancing activities in serum samples which were collected from dengue patients at multiple time points between the acute and defervescent phases (Figs. 1 and 2). Patient #46 showed DHF manifestations in spite of the strong neutralization against both autologous and prototype viruses, and the dose-dependent curve was not meaningfully shifted over 0–19 h in this patient. This suggests that the strong neutralizing antibodies that had been induced by four days after the fever onset might not have protected the patient from the subsequent severe disease, although the antibody status in the pre-infection phase was not known. In contrast, DF patient #44 did not show severe manifestations, even though the ADE activity patterns of this patient were as high as those of the DHF patients (#57 and #49). Therefore, when samples from post-hospitalized patients were used, it was still difficult to predict disease outcomes by measuring the balance between neutralizing and enhancing activities. Although other factors might be involved in determining the immune correlation (such as the treatment conditions in the hospital, current infecting serotype and previous infection history, etc.), the most valuable factor (and also the most difficult) is the collection of samples in the pre-infection or pre-hospitalization period through prospective cohort studies (22-25).

The ADE activities shown by the low serum dilution (1:10) were gradually decreased over time (Fig 3B and Supplemental Fig. S1). These results suggest that neutralizing antibodies were immediately produced after the infection, and then the original enhancing activity might have been competitively suppressed by the appearance of neutralizing antibodies. However, some cases did not show a decreasing trend in the heterologous combinations (Supplemental Fig. S1). For instance, patient #44 displayed the decreasing trend against DENN-2 (autologous and NGC strains) and DENV-4, but exhibited an increasing or flat trend against DENV-1 and DENV-3. This suggests that patient #44 will be protected from subsequent infection with DENV-2 and DENV-4, but may not be protected from infection with DENV-1 or DENV-3. Such patients should be advised to take care to avoid future infections. Therefore, monitoring the balance between NAb and EAb activities against all serotypes is important for the prediction of disease severity upon secondary heterotypic infection in seropositive individuals. On the other hand, the opposite trend was observed in the high-serum dilution range (1:2560)—that is, the ADE activities were basically increased over time (Supplemental Fig. S2). This suggests that the induced neutralizing antibodies still possess the potential ADE activity at sub-neutralizing doses. Therefore, if the antibody level wanes over time, it might become a risk factor for increased disease severity.

The first approved dengue vaccine, Sanofi’s Dengvaxia, has been introduced into approximately 20 endemic countries since 2015 (26). However, WHO SAGE revised their recommendation to say that only seropositive individuals, who have previously been infected with DENV, can be inoculated with Dengvaxia (27). Seronegative populations should not be vaccinated (28, 29), because a risk of vaccine-induced infection enhancement cannot be excluded (30-32). Although there are several assay kits to detect dengue antibodies in a quantitative or qualitative manner (e.g., ELISA, immunochromatography assays, etc.), few rapid test kits are available to detect the functional antibody activity. Only an assay with the capacity to detect ADE activity would be able to provide a risk assessment for vaccine recipients. In the present study, five (#44, #46, #57, #12 and #49) of six patients were confirmed to be seropositive (classified as secondary infection). In contrast, patient #14 (classified as primary infection) showed no functional antibody activity against the autologous DENV-1, NGC (DENV-2) and H241 (DENV-4) strains (Figs. 1 and 2), although this patient did show limited enhancing activities against the Mochizuki (DENV-1) and H87 (DENV-3) strains. Thus, such individuals should be advised to delay vaccination until their antibody levels are elevated sufficiently, if they have a plan to get Dengvaxia in the near future.

A previous study showed that ADE contributed to an increase in viremia in an animal model (33), and the high viremia levels in dengue patients have been demonstrated to correlate with the disease severity (6, 9, 10). Since the viremia in humans is caused by secretion of the virus from the infected-host cells, the *in vitro* progeny virus infectivity obtained from the NAb/EAb-balance assay might correlate to the clinical viral load. Interestingly, in the present study, significant positive correlations were observed between *in vitro* ADE progeny virus levels (focus forming units [FFU]/ml) in serum samples and *in vivo* viremia levels (copies/ml) in plasma samples (Fig. 4B). This result suggests that measurement of the infective progeny virus titers (or the infected cell counts) in the NAb/EAb-balance assay may enable us to predict the impact of viral load in the subsequent DENV infection. Since patients #14 and #46 showed no enhancing activity against autologous virus in the lower serum dilutions (Fig. 1B), the viral load quantity in the plasma might have been lower in these patients than the other participants (Fig. 4A).

In conclusion, the present NAb/EAb-balance assay might be used to predict the impact of the viremia level when a human is subsequently infected with DENV.

Although we could not find a relationship between the ADE activity (viremia level) in the early phase and disease severity in the defervescent phase, the *in vitro* progeny virus titers obtained from the NAb/EAb-balance assay were shown to significantly correlate with the viral load determined by qPCR assay. The major limitation of this study was the small sample size. Nonetheless, although only six autologous virus strains were successfully isolated in this study, we believe that the analysis of the combination of autologous virus and serum is meaningful to understand the relationship between neutralization/ADE and disease outcome.

## MATERIALS AND METHODS

### Blood samples

The present study was conducted using the serum and plasma collected from confirmed-dengue patients at the Hospital for Tropical Diseases, Bangkok, Thailand, in a previous study (21). Six dengue patients were enrolled from whom clinical virus isolates were obtained. All subjects gave their informed consent for inclusion into the study before they participated. This study was conducted in accordance with the Declaration of Helsinki, and the protocol was approved by the Ethics Committee of the Faulty of Tropical Medicine, Mahidol University, Thailand (FTM ECF-019-04).

### Cells

African green monkey kidney Vero cells were cultivated in Eagle’s minimum essential medium supplemented with 10% fetal bovine serum (FBS) and 60 μg/mL kanamycin (34). Human erythroleukemia K562 cells were cultivated in RPMI 1640 medium supplemented with 10% FBS, 100 units/mL penicillin, and 100 μg/mL streptomycin (35). All cell lines were cultivated in a humidified atmosphere of 5% CO_2_:95% air at 37 °C.

### Virus isolation and typing

Vero cells were inoculated with the diluted patient sera (1:10) collected in the acute phase, and then incubated at 37 °C for 7 days. Within five blind passages, viral RNA was extracted from the supernatant of the infected cells, and the serotype was determined by PCR using type-specific primers following a previous report (36).

### Viruses

Six clinical isolates (1 strain of DENV-1, 2 of DENV-2, 1 of DENV-3, and 2 of DENV-4) and four prototype lab strains (DENV-1: Mochizuki strain; DENV-2: New Guinea C [NGC] strain; DENV-3: H87 strain; and DENV-4: H241 strain) (37) were used in this study. The culture supernatants harvested from the infected Vero cells were used as live virus sources for the antibody assay measuring the balance between neutralizing and enhancing activities (see below).

### Antibody assay for the balance between neutralizing and enhancing activities (NAb/EAb-balance assay)

The NAb/EAb-balance assay was conducted using semi-adherent K562 cells as described previously (19). Briefly, serial dilutions of sera (starting from 1:10 dilution) were mixed with each DENV strain in a poly-L lysine-coated 96-well microplate and incubated at 37 °C for 2 h. K562 cells (1 × 10^5^ cells per well) were then added to the mixtures and incubated at 37 °C for 2 days. After immunostaining (see below), the infected cells were counted. The cut-off values for neutralizing and enhancing activities were calculated from the means ± three standard deviations (SD) of infected cell counts obtained with eight negative controls adjusted for approximately 100 infected cells. When the number of infected cells was higher than the mean + three SD, it was defined as enhancing activity. In contrast, when the infected cell number was lower than the mean - three SD, it was defined as neutralizing activity.

In addition, enhancing activity was also evaluated by titrating the progeny virus infectivity (FFU/ml) in the culture supernatant of the infected cells. The supernatants were harvested at 24 h after the inoculation, and their titers were determined on Vero cells.

### Immunostaining

Immunochemical staining was performed essentially as described previously (34). Briefly, cells were fixed with acetone/methanol (1:1) and incubated serially with a mouse monoclonal antibody D1-4G2 (flavivirus group cross-reactive) purchased from American Type Culture Collection (Manassas, VA), biotinylated anti-mouse IgG, ABC (avidin-biotinylated peroxidase complex) reagent, and VIP substrate (Vector Laboratories, Burlingame, CA).

### Quantification of the viral RNA copy number in plasma samples

The viral RNA copy number in plasma samples was determined by following a previous study (21). Briefly, viral RNA was extracted from 70 μl plasma using a QIAamp Viral RNA Mini Kit (Qiagen, Hilden, Germany) according to the manufacturer’s protocol, then subjected to quantitative real-time RT-PCR using a One-Step SYBR PrimeScript RT-PCR kit II (Takara Bio, Japan). The PCR mixture was mixed with 2 μl of extracted RNA and DENV-specific primer (38) before running on a CFX96TM real-time PCR cycler (BioRad, Hercules, CA, USA) under cycle conditions of 42°C for 5 min, 95°C for 10 sec followed by 45 cycles of 95°C for 5 sec, 55°C for 30 sec and 72°C for 30 sec. The viral load quantity was determined by linear regression of the cycle threshold value against the known viral titers quantified by plaque forming unit assay.

### Statistical analysis

Correlation coefficients were estimated on the basis of the Pearson product-moment correlation coefficient (r). A probability (*p*) less than 0.05 was considered statistically significant.

## Acknowledgments

This research was supported by the Japan Initiative for Global Research Network on Infectious Diseases (J-GRID) from the Ministry of Education, Culture, Sport, Science & Technology in Japan, and the Japan Agency for Medical Research and Development (AMED) (JP18fm0108003).

## Supplemental Figure Legend

**Supplemental FIG S1. Transition of the NAb/EAb balance activity in a low serum dilution (1:10) with time progression.** The infected cell counts (specifying data at 1:10 serum dilution) obtained from Figs. 1B and 2 were plotted as the ordinate (expressed as log10) against time (h) as the abscissa.

**Supplemental FIG S2. Transition of the NAb/EAb balance activity in a high serum dilution (1:2560) with time progression.** The infected cell counts (specifying data at 1:2560 serum dilution) obtained from Figs. 1B and 2 were plotted as the ordinate (expressed as log10), and plotted with time (h) as the abscissa.

